# Genetically distinct behavioral modules underlie natural variation in thermal performance curves

**DOI:** 10.1101/523654

**Authors:** Gregory W. Stegeman, Scott E. Baird, William S. Ryu, Asher D. Cutter

## Abstract

Thermal reaction norms pervade organismal traits as stereotyped responses to temperature, a fundamental environmental input into sensory and physiological systems. Locomotory behavior represents an especially plastic read-out of animal response, with its dynamic dependence on environmental stimuli presenting a challenge for analysis and for understanding the genomic architecture of heritable variation. Here we characterize behavioral reaction norms as thermal performance curves for the nematode *Caenorhabditis briggsae*, using a collection of 23 wild isolate genotypes and 153 recombinant inbred lines to quantify the extent of genetic and plastic variation in locomotory behavior to temperature changes. By reducing the dimensionality of the multivariate phenotypic response with a function-valued trait framework, we identified genetically distinct behavioral modules that contribute to the heritable variation in the emergent overall behavioral thermal performance curve. Quantitative trait locus mapping isolated regions on Chromosome II associated with locomotory activity at benign temperatures and Chromosome V loci related to distinct aspects of sensitivity to high temperatures, with each quantitative trait locus explaining up to 28% of trait variation. These findings highlight how behavioral responses to environmental inputs as thermal reaction norms can evolve through independent changes to genetically distinct modular components of such complex phenotypes.

**Article Summary:** Plastic responses to environmental inputs, reaction norm phenotypes that can be summarized with parameters of fits to a mathematical function, are pervasive across diverse organismal traits and crucial to organismal fitness. We quantified the nematode *Caenorhabditis briggsae*’s behavioral thermal performance curves as function-valued traits for 23 wild isolate genotypes and 153 recombinant inbred lines. We identified quantitative trait loci on multiple chromosomes that define genetically distinct behavioral modules contributing to the emergent overall behavioral thermal performance curve. These findings highlight how dynamic behavioral responses to environmental inputs can evolve through independent changes to genetically distinct modular components of such complex phenotypes.

## Introduction

Plastic responses to environmental inputs, reaction norm phenotypes, are pervasive across diverse organismal traits and crucial to organismal fitness (Schlichting and Pigliucci 1998; Stinchcombe and Kirkpatrick 2012). Such phenotypic plasticity can manifest as irreversible switches during development, physiological or metabolic adjustments, or rapid behavioral reactions (Dingemanse *et al.* 2010). This ability to vary phenotype in response to environmental conditions can be adaptive, though maintaining the biological architecture required for adaptive plasticity may be costly or evolutionarily constrained (Murren *et al.* 2015; Nettle and Bateson 2015). Like any trait, reaction norm phenotypes can vary genetically in the wild so that selection may act on the genetic variation to shape the magnitude and form of phenotypic plasticity (Stinchcombe and Kirkpatrick 2012). Two outstanding key questions in the evolution of plastic traits include how genetically complex is the control of natural variation in such seemingly complex phenotypes and how genetically modular are they.

Thermal performance curves (TPCs) represent a common type of reaction norm in which a trait depends on temperature conditions in a stereotypical way (Huey and Stevenson 1979; Huey and Kingsolver 1989). Temperature is an ever-present environmental factor that strongly modulates diverse aspects of biology, especially traits of ectothermic organisms (Fischer and Karl 2010; Colinet and Hoffmann 2012). Ectotherms have evolved diverse physiological strategies and molecular mechanisms to adjust to changes in temperature (Fields 2001). Behavior, however, remains the principal means by which ectothermic animals regulate body temperature by sensing, orienting and navigating their thermal landscape in ways crucial for fitness (Huey and Kingsolver 1993; Garrity 2007). Behaviors are notoriously plastic and dynamic traits controlled by networks of many genes that thus lend themselves to quantitative genetics approaches to relate genotype and phenotype (Anholt and Mackay 2004; Dingemanse *et al.* 2010).

Summarizing the complexity of such dynamic traits presents a challenge in the analysis and understanding of plastic reaction norm phenotypes (Stinchcombe and Kirkpatrick 2012). One way to simplify them is to describe the functional form of the trait response to environmental changes with fits of appropriate mathematical equations to experimental data. The corresponding parameter estimates of the function can then be treated as phenotypes themselves, termed ‘function-valued traits’ (Ma *et al.* 2002; Wu and Lin 2006; Yap *et al.* 2007; Stinchcombe and Kirkpatrick 2012). Functional forms with biologically intuitive interpretations of the parameters are especially appealing for distilling multidimensional phenotypes into a few key, distinct components. Partitioning complex phenotypes as function-valued traits in this way also is powerful for mapping trait variation to genomic variation with approaches like quantitative trait loci (QTL) mapping (Baker *et al.* 2015). Using function-valued traits instead of a single trait or a long list of univariate traits as phenotypes for QTL mapping may provide extra biological insight and statistical power to detect the most relevant QTLs affecting a reaction norm phenotype (Wu and Lin 2006; Stinchcombe and Kirkpatrick 2012).

Analysis of behavioral reaction norms in *Caenorhabditis elegans* demonstrated a neural ‘decision-making’ contribution to temperature-dependent locomotory responses using the same experimental paradigm as we here investigate in its congener *C. briggsae* (Stegeman *et al.* 2019). *C. briggsae* may provide an even more powerful animal model system to investigate natural genetic variation in temperature-dependent behavioral reaction norms. In contrast to *C. elegans*, *C. briggsae* displays latitudinal genetic differentiation and adaptive differentiation to distinct thermal regimes (Cutter 2015), with genotypes from the “Tropical” phylogeographic group having higher fitness at extreme warm temperatures and the “Temperate” group genotypes more fecund at extreme low temperatures (Prasad *et al.* 2011). While the vast majority of isolates of *C. briggsae* found around the world fall into these two phylogeographic groups, five other genetically distinctive groups of restricted geographic origin also are known to contribute genetic variability to the species (Cutter *et al.* 2006; Felix *et al.* 2013; Thomas *et al.* 2015). These distinctive genetic backgrounds also show temperature dependent behavior differences (Stegeman *et al.* 2013). Moreover, the high-quality genome assembly and annotation for *C. briggsae* and a library of advanced-intercross recombinant inbred lines (RILs) with dense genotype information provides a powerful experimental stage for connecting phenotypic differences to natural genetic differences (Ross *et al.* 2011). This context prompted us to investigate temperature-dependent behavioral reaction norms among natural isolate strains and in genetic mapping populations of *C. briggsae* with the aim of providing insight into the genetics and evolution of plastic behavioral traits.

Here we quantify behavioral reaction norms as thermal performance curves for locomotion in response to increasing or decreasing temperature. Quantitatively, one may distill the complex shape of TPCs into characteristic parameters that describe the functional form of the response, treating the emergent multidimensional phenotype as a set of function-valued traits (Stinchcombe et al., 2012). We found that TPC variation induced by different environments can be dissociated into distinct genetic components. We demonstrate substantial natural variation in TPCs among wild genetic backgrounds in addition to “classic” developmental plasticity in TPC shape in response to distinct rearing conditions. By quantifying variation in behavioral reaction norms for F1 heterozygous lines, recombinant inbred lines, and introgression lines, we map distinct QTL for components of the TPCs as function-valued traits to show that genetically separable phenotypic modules underlie the genetic architecture of the overall emergent trait captured with behavioral thermal reaction norms.

## Methods

### Experimental quantification natural genetic variation in locomotory behavior

We quantified nematode locomotory behavior as a function of changes in temperature using the liquid micro-droplet experimental paradigm that we developed previously (Stegeman *et al.* 2019). In brief, we recorded video and quantified the swimming behavior of individual worms placed into 1.8μL droplets arranged in an array of 18 droplets on a glass slide as we manipulated temperature with custom hardware (Stegeman *et al.* 2019). Image capture and processing of swimming worms was performed with custom programs written in LabVIEW (National Instruments, Austin, TX) with further data analysis in R to compute metrics of locomotory activity (Stegeman *et al.* 2019). Our standard assay involved an increasing temperature regime from 25°C to 41°C with 1°C steps occurring every 80 s, recording video of the behavior for the last 20 s of each temperature step at 15fps. We also performed experiments on wild isolates with a decreasing temperature regime (from 25°C to 8°C with 1°C steps).

To investigate variation in TPCs, we reared 23 *C. briggsae* natural isolates at 16°C, 23°C and 28°C to quantify genetic and plastic contributions to locomotory behavior profiles in the liquid micro-droplet assay when we exposed animals to increasing or decreasing ambient temperature changes from an initial 25°C (Supplementary Table S1). The fitness optimum for the Tropical AF16 strain is close to 21.5°C, with 16°C and 28°C both yielding strongly reduced fitness (Prasad *et al.* 2011; Begasse *et al.* 2015). These strains derive from genetically distinct phylogeographic origins known to exhibit phenotypic variation for temperature-related traits (Prasad *et al.* 2011; Stegeman *et al.* 2013), with 16 of them having published genome sequence data (Thomas *et al.* 2015). We also generated and tested reciprocal F1 heterozygotes from crosses of two strains (Tropical AF16 and Temperate HK104), using parental hermaphrodites that were depleted of self-sperm to ensure harvesting of non-selfed F1 offspring. Experimental assays included only well-fed adult worms with *E. coli* OP50 from non-starved NGM agar plates kept at 23°C unless indicated otherwise.

For analysis of wild isolates, we tested 36 worms from each strain from each rearing temperature, with each block of 18 droplets containing 9 replicate worms of two strains. We randomized testing order combinations of the 23 wild isolate strains over a period of a week separately for the increasing and decreasing temperature assays.

For QTL mapping, we phenotyped a panel of 153 recombinant inbred lines (RIL) that are genomic mosaics derived from AF16 and HK104 parental lines (Ross *et al.* 2011). These 153 lines were genotyped at 1031 single nucleotide polymorphism (SNP) markers across the six chromosomes, which define 430 distinct genetic blocks for this subset of the full RIL collection created by (Ross *et al.* 2011) (Supplementary Table S2). We performed our standard liquid micro-droplet assay on the RIL and parental strains with ~1°C temperature increments from 25° to 41°C for animals reared at 23°C. We tested 45 worms from each homozygous RIL strain in a randomized order with 9 replicate animals for each of two strains on each assay slide with additional replicates for the parental lines AF16 and HK104. After quality filtering, we included data from 6907 individual animals across the 155 strains, with data from ≥40 animals for each strain (except PB1209 with *n*=36). The number of animals that passed quality control at a given temperature point can be lower than 40 to still be included in analyses, provided ≥25 replicate animals were of acceptable quality for analysis (Supplementary Figure S1).

We calculated several distinct but inter-related metrics of worm movement behavior: locomotion index with 1/15^th^ second frame resolution (LI1), locomotion index with 7/15^th^ second frame resolution (LI7), pixel center-of-mass displacement (at 1/15^th^ and 7/15^th^ second frame resolution), and the proportion of worms moving at each temperature step (calculated from a threshold of activity for LI7). We calculated the LI index of locomotion for each individual animal at each temperature step as the fraction of overlapping pixels of the worm between different frames of video: adjacent video frames 1/15th second apart (LI1) or 7/15^th^ second apart (LI7). The LI values range from 0 (no movement detected) to a maximum of 1. We standardized the raw LI1 and LI7 values for each individual’s locomotion by subtracting its lowest value across temperature steps to account for the slight pixel noise across video frames even for immobile animals. The proportion of animals moving provides an overall indicator of activity across replicate individuals, calculated after we assigned values of 1 (movement detected with LI7) or 0 (no movement detected with LI7) for each individual worm from which we computed the mean and standard deviation at each temperature for each strain. These locomotion metrics are not fully independent and their individual values will correlate between adjacent temperature steps. Consequently, we sought to reduce the phenotypic dimensionality into independent components using two techniques: function fitting and principal component analysis.

### Behavioral thermal reaction norms as function-valued traits

In order to compare thermal performance curves as a whole, rather than considering separately the metrics at each temperature, we fit a three-parameter logistic function to the behavioral thermal response curve from benign to extreme temperatures for each strain as:

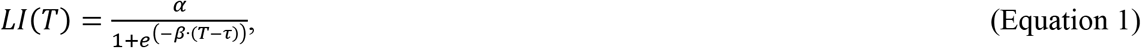

where *T* is droplet temperature, *α* is the asymptote that describes the baseline locomotion at the beginning of the assay (performance at temperatures near 25°C), *β* is the slope parameter that corresponds to how rapidly locomotion declines as temperature rises or falls, and *τ* is the inflection point parameter, which represents the temperature at which the thermal performance curve switches from concave to convex (Figure 1). Prior to function fitting, we normalized each individual’s locomotion by subtracting its lowest LI1 (or LI7) value (usually ~0.06-0.12) from the scores at all other temperature steps to account for the slight pixel noise across video frames when worms are immobile that keep LI scores from reaching 0. We performed functional fits to mean LI1 or LI7 values from each temperature step from a given treatment by using the three parameter logistic equation in the non-linear model fitting procedure in JMP v10.0.0 (SAS Institute). This procedure could capture most of the change in strain mean locomotion scores at each temperature (*R*^2^ of 0.98-1.00). We compared pairs of strains by treating TPCs as function-valued traits in this way and used the 95% confidence intervals for the parameter estimates to determine whether pairs of strains differed significantly from one another.

**Figure 1.**
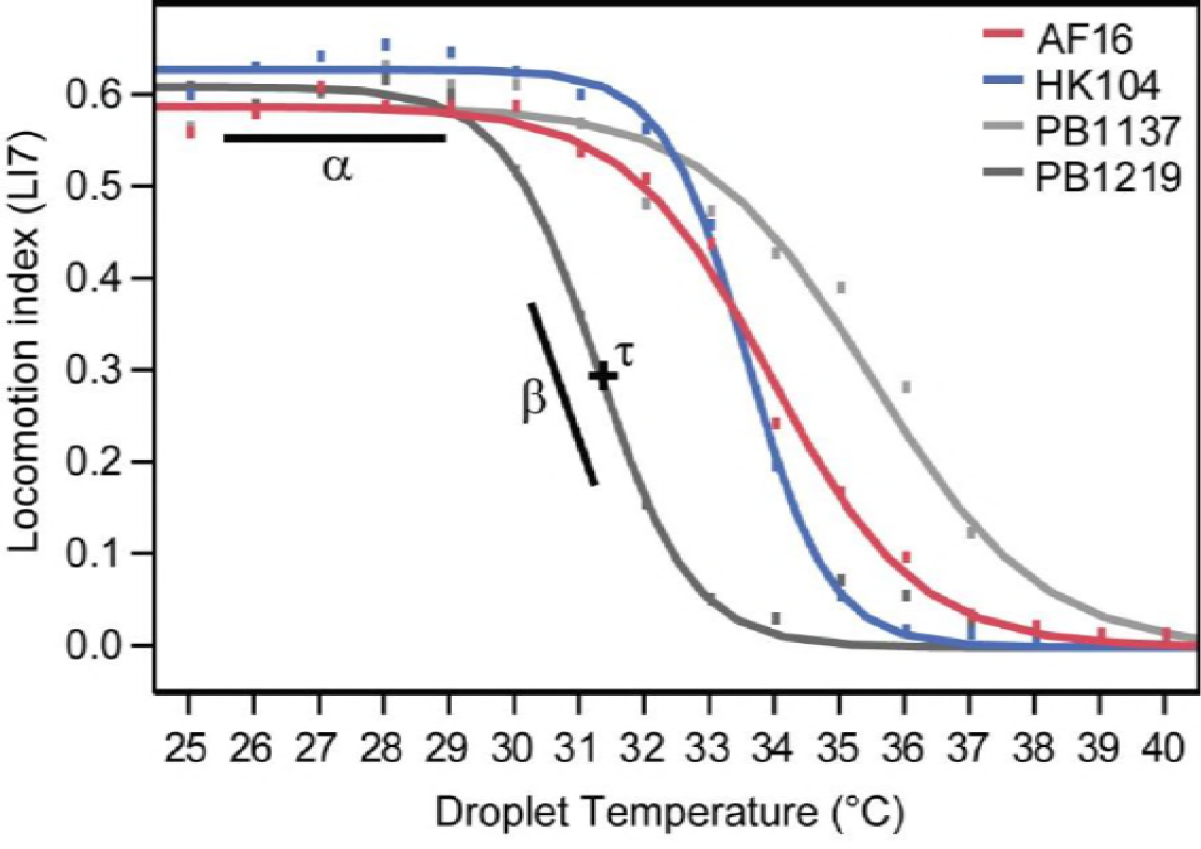
Three parameter logistic function fit for four example strains, including the RIL parents that correspond to Tropical and Temperate phylogeographic groups (AF16, HK104) and two RILs with extreme TPCs (PB1137, PB1219). Locomotion index (LI7) adjusted by subtracting each worm’s minimum score from the value at each temperature step prior to fitting strain mean values to Equation 1.

Many analyses of thermal performance focus on thermal optima, or peak performance, which is not captured in the logistic function fits described above. In order to estimate TPC peak performance for natural isolates strains, we fit a Gaussian peak function (Equation 2) to a subset of locomotion data. The TPC at relatively benign temperatures between 18°C to 32°C exhibits minimal skew, so we fit mean values combined for individuals from increasing and decreasing temperature assays with the non-linear model fitting procedure in JMP v10.0.0 (SAS Institute):

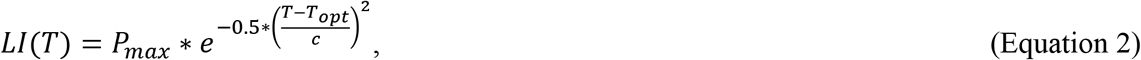

where *T* is droplet temperature, *P*_max_ is the peak value for the locomotion index, *T*_opt_ is the ‘critical point’ temperature where the peak occurs, and *c* is the curvature.

### Behavioral dimensionality reduction with Principal Component Analysis

As an alternate means of reducing the dimensionality of the data by integrating correlated phenotype metrics, we used principal components analysis (PCA) on the phenotypic measurements of the 153 RIL strains. We performed PCA using the rda (redundancy analysis) function with scaling enabled, from the vegan package (v2.3-3) in R (with strain as “sites” and phenotypes as “species”). We included univariate phenotypes for the strain mean and standard deviation at each temperature step for five metrics of locomotory activity (LI1, LI7, 1 and 7 frame displacements, and proportion moving). We also included the mean pixel area for each strain as well as the three parameter logarithm fit parameter estimates and standard errors for a total of 167 phenotypic metrics for each strain.

### Genetic mapping of natural variation in locomotory behavior

We used standard interval mapping to associate genotype and phenotype of RIL strains, as implemented in the r/qtl package version 1.39-5 (Broman *et al.* 2003) in R v. 3.2.3 (R Core Team, 2015). To fill in potential break points between genotyped markers, we applied the multiple imputation method for standard interval mapping in this package. Specifically, we simulated 128 possible combinations of break points at 0.5cM spacing between genotyped markers. For the two-dimensional QTL analyses, we used Haley-Knott regression instead of multiple imputations in order to reduce the computational load.

We first QTL mapped the 8 ‘synthetic’ principal component locomotion behavior traits using 1031 markers across the six chromosomes plus two pseudo-markers representing the mitochondrial genome. We determined a 5% genome wide significance threshold for each trait from 2500 permutations and Bonferroni correction (P=0.05/8). We followed a similar procedure in QTL mapping the three function fit parameter phenotypes for locomotion index (normalized LI7), using significance thresholds derived from 2500 permutations of the data with Bonferroni corrected P=0.05/12. We then computed 95% Bayes credible intervals for log-odds (LOD) peaks on chromosomes with significant QTL. We estimated the broad-sense heritability of QTL at the peak marker location as 1 – 10^−2 LOD/*n*^ (*n* = number of RILs). We also performed two-dimensional QTL scans in r/qtl with a minimum distance of at least 5cM between markers and 2500 permutations of the data to estimate significance for a subset of phenotypes. Model type was determined as an interaction if the full model allowing for interactions (lod.fv1) was significant while the additive model (lod.av1) was not; the model type was resolved as additive if the lod.av1 model was significant, and lod.fv1 was not, or if the interaction between these (lod.int) was not significant (Broman and Sen 2009).

The collection of *C. briggae* RIL strains exhibited marker transmission ratio distortion (Ross *et al.* 2011). These included significant bias towards HK104 markers on Chromosome V near the QTL peak that we identified for high temperature locomotion. There were also allelic biases on Chr III and Chr IV (Ross *et al.* 2011). This likely contributes to fewer novel breakpoints in the NILs for these regions than expected from cross-over rate estimates. While some inadvertent selection due to slower development during the intercrossing of the RILs, it is primarily associated with distortion on chromosome III (Ross *et al.* 2011). The distortions on Chromosomes V and IV are only significant for one of the initial F1 cross directions, and show no LD with the X chromosome, which suggests a cytonuclear epistatic interaction (Ross *et al.* 2011; Chang *et al.* 2015).

To investigate the QTL region on Chromosome V, we generated 13 near isogenic lines (NILs) by backcrossing RIL strains PB1124 and PB1147 to AF16. We used restriction-digest fragment length polymorphism (RFLP) marker-assisted selection of backcross individuals for genotyping near recombination breakpoints in the QTL region of interest through at least 7 generations of backcross. After selfing the resulting backcross lines, we then affirmed homozygosity and genotype along Chromosome V, as well as for three loci for each of the other chromosomes, using markers in (Koboldt *et al.* 2010) (loci with “cb#” SNP names in Supplementary Table S3). We also designed and used four additional RFLP markers to increase marker density along Chromosome V (Supplementary Table S3). We tested these lines using the same droplet assay with increasing temperatures as we used for the RIL QTL analysis. We also tested the NILs and parental lines in modified droplet assays to explore alternative related phenotypes.

We tested for gene ontology (GO) term enrichment within the QTL region on Chromosome V using PANTHER (http://www.pantherdb.org/), which we visualized using Treemaps created with REVIGO (Supek *et al.* 2011); 33 of the 115 genes lacked GO terms.

### Data and reagent availability

All *C. briggsae* strains are available from the Caenorhabditis Genetics Collection (AF16, HK104, wild isolates) or from the authors (recombinant inbred lines, near isogenic lines). Supplementary tables, figures and data are available in FigShare. File S1 contains Supplementary Tables S1 and S3-S6 as well as Supplementary Figures S1-S12. File S2 contains Supplementary Table S2 (genotypes for each RIL used in QTL mapping) and File S3 contains Supplementary Table S3 (genes with coding changes and GO terms). Per-individual phenotype data is given in Files S4, S5, S6, and S7 for F1s, wild isolates, recombinant inbred lines, and near isogenic lines, respectively. Per-strain phenotypes of RILs used in QTL mapping for all univariate traits are provided in File S8.

## Results

We hypothesized that rearing temperature would differentially affect thermal performance of swimming behavior of distinct genotypes, motivated by prior studies that demonstrated temperature-dependent trait differences among phylogeographic groups of *C. briggsae* (Prasad *et al.* 2011; Stegeman *et al.* 2013). We therefore quantified natural genetic variation and environmental plasticity in locomotion behavior in *C. briggsae* by assaying thermal performance with a liquid micro-droplet experimental paradigm described previously (Stegeman *et al.* 2019).

In these experiments, we isolated individuals in ~2μl liquid droplets arranged in an array on a microscope slide and incremented their temperature from a benign starting point (25°C) in ~1°C steps to hot or to cold extremes (41°C or 8°C) while taking video recordings for automated image analysis and behavioral quantification. Like diverse other organisms (Huey and Stevenson 1979; Huey and Kingsolver 1989), including *C. elegans* (Begasse *et al.* 2015; Stegeman *et al.* 2019), *C. briggsae* generally reacts to changing temperature with a functional response like a classic thermal performance curve (TPC): peak activity at intermediate temperatures, with a shallow rate of declining locomotion toward cold temperatures, and a rapid reduction in movement toward high temperatures (Figure 2). We then quantified the behavioral TPCs as function-valued traits to reduce the dimensionality of the phenotype and extract the dynamic behavioral information into biologically-meaningful parameters of the function fits (Figure 1). Thermal performance curves for 23 natural isolate strains from seven genetically distinct phylogeographic groups of *C. briggsae* (Table 1; (Cutter *et al.* 2006; Felix *et al.* 2013)) revealed substantial heritable and environmental contributions to variation in behavioral TPC shape (Figure 2, Figure 3). By then focusing on 153 recombinant inbred lines (RIL) derived from the Tropical and Temperate phylogeographic groups to investigate the genetic differences underlying variation in thermal performance, our quantitative trait loci (QTL) mapping identified genetically distinct behavioral modules that contribute to TPC shape.

**Figure 2.**
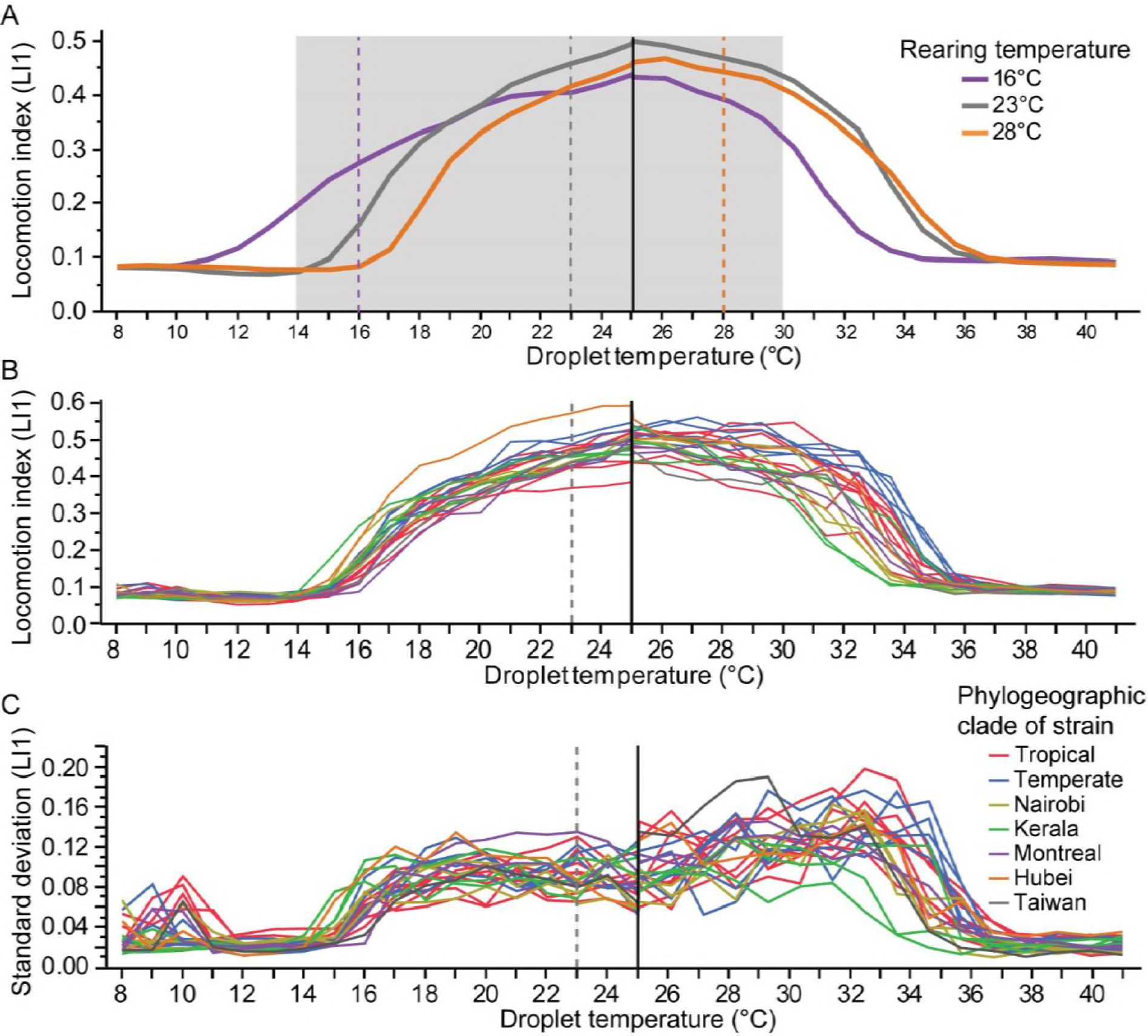
Thermal performance of *C. briggsae* reared at three growth temperatures (purple=16°, grey=23°C, orange=28°C), with data pooled across experiments for the raw LI1 locomotion index (**A**). Assays either incremented or decremented temperature from a 25°C initial assay temperature (black vertical line). (**B**) TPCs for each of the 23 wild isolate strains reared at 23°C (raw LI1; ~36 worms per strain), colored by phylogeographic group (8 Tropical red; 5 Temperate blue; 3 Nairobi yellow; 3 Kerala green; 2 Montreal purple; 1 Hubei orange; 1 Taiwan black). (**C**) Standard deviation of locomotion index values (raw LI1) for each wild isolate strain shown in (B). Rearing temperatures are denoted with vertical dashed lines; gray region in (A) indicates fertile temperature range. The spikes in SD and LI at temperatures below 12°C are due to condensation artifacts in the image analysis.

**Figure 3.**
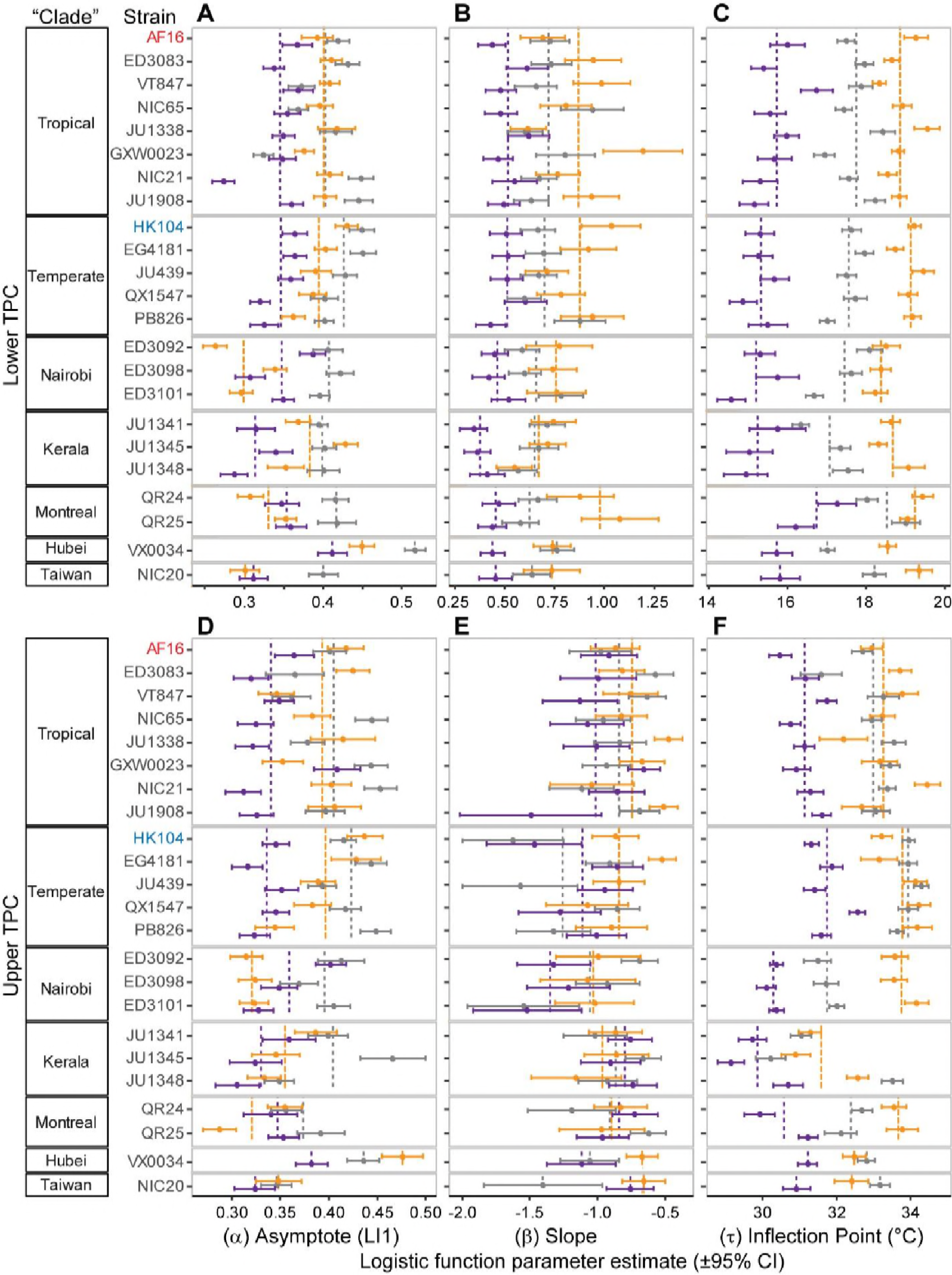
Parameter estimates from logistic function fits to locomotion index (LI1) thermal performance curves for each wild isolate strain (see Equation 1 and Figure 1). Parameter values for decreasing temperature TPCs (**A, B, C**) and increasing temperature TPCs (**D, E, F**) estimated separately for animals reared at each of three temperatures (purple=16°C, gray=23°C, orange=28°C). Dashed lines indicate the mean across strains within a phylogeographic group (colored by rearing temperature); error bars indicate ±95% confidence interval.

### Thermal performance curves are sensitive to genotype and environment

Under all three rearing temperature conditions (16°C, 23°C and 28°C), the TPCs for locomotion are unimodal with declines in performance towards low and high temperatures. For animals reared at cool to moderate temperatures, the TPC shows the classic shape of a steeper drop in performance toward high than low temperatures, but those reared at a high 28°C actually tend to show sharper performance declines toward low temperatures (Figure 2, Figure 3; Supplementary Figure S2). Thermal performance curves also shift position and width depending on rearing temperature, with cooler rearing temperatures yielding wider TPCs that show more locomotory activity at lower temperatures (Figure 2, Figure 3; Supplementary Figure S2).

Despite rearing temperature yielding large and consistent differences in locomotion that give all genotypes a similar characteristic TPC shape (Ayrinhac *et al.* 2004; Hoffmann *et al.* 2005), the genetic identity of strains also contributes substantial variation in temperature-dependent locomotion profiles (Figure 2). While strain-specific genotypic differences appear more pronounced than general differences among phylogeographic groups, visual inspection of TPCs for the genotypes from the Kerala “clade” strains exhibit cold-shifted TPCs compared to most other strains (Figure 2, Figure 3). By contrast, the two Montreal “clade” genotypes tend to perform poorly at colder temperatures and the Hubei genotype has the highest and one of the broadest TPCs (Figure 3).

To quantify these trends, we applied a 3-parameter logistic function (Equation 1) to data from each half of the TPC (the ‘cool’ half with subscript “C” and the ‘hot’ half with subscript “H”). From these function fits to data, we extracted the fit parameter values as simple and biologically interpretable summaries of locomotion, treating the reaction norm phenotype as a function-valued trait to compare TPCs across strain and rearing condition treatments.

First, we characterized baseline swimming behavior at benign temperatures with the asymptote parameter (*α*), which is related to maximum locomotion. We observed *α* to be highest for most strains when animals were raised at a benign 23°C and *α* lowest when raised at 16°C (ANOVA *α*_C_ F_2,66_=18.48, P<0.0001; *α*_H_ F_2,66_=15.57, P<0.0001; Tukey’s posthoc tests show differences among all rearing temperatures for both *α*_C_ and *α*_H_) (Figure 2, Figure 3, Supplementary Table S4). Animals reared at 23°C appear visibly healthier than when reared at the extreme temperatures, and this may translate into them being capable of faster locomotion at initial assay temperatures. Alternatively, the thermal jump from rearing temperature to the initial assay temperature (25°C) is largest for animals reared at 16°C, which might shock them into slow initial locomotion as captured by *α*. We also observed significant genetic variation in both *α*_H_ and *α*_C_ (non-overlapping 95% CIs among genotypes; Figure 3). The Hubei strain VX34 shows consistently fast *α* across rearing temperatures, whereas the three strains from Nairobi consistently exhibit especially slow baseline locomotion when reared at 28°C (Figure 3). Strain genotypes also differ considerably in their degree of plasticity to rearing conditions, with some strains having nearly identical *α*_C_ values regardless of rearing temperature (overlapping 95% CI across rearing temperatures for e.g. VT847, NIC65, GXW23) and others being very sensitive (non-overlapping 95% CI across rearing temperatures for e.g. NIC21, JU1348, VX34; Figure 3). This variation in trait canalization also extends to the hot end of the TPC (e.g. compare low vs high plasticity in *α*_H_ for VT847, QR24, NIC20 vs ED3083, NIC21, VX34; Figure 3). These strain differences in plasticity point to genetic variation for the phenotypic robustness of baseline locomotion behavior in response to rearing conditions, in addition to heritable variation for the raw locomotory response.

Our function fit captures the sensitivity to changes in ambient temperature as the sharpness of the decline in locomotory activity, with this slope parameter (*β*) also influenced by rearing temperature and genotype (ANOVA *β*_C_ F_2,66_=55.94, P<0.0001; *β*_H_ F_2,66_=3.75, P=0.029; non-overlapping 95% CIs among genotypes) (Figure 2). The magnitude of the slope (|*β*|) is steeper for most strains when the assay increments toward hot temperatures than when it approaches cool temperatures, at least when they were reared under cool or benign conditions (Figure 3, Supplementary Table S4). For animals derived from these same cool or benign rearing conditions, we also observe more genetic variation in *β* when they experience the warming portion of the TPC compared to cooling half of the TPC (F-test for variances: 16°C rearing *F*_22,22_=0.137, *P*<0.0001; 23°C rearing *F*_22,22_=0.097, *P*<0.0001; 28°C rearing *F*_22,22_=0.828, *P*=0.66). Moreover, the cooler the rearing temperature, the shallower the slope for the decline in locomotion toward low temperatures (Tukey posthoc tests indicates *β*_C 16°_ < *β*_C 23°_ < *β*_C 28°_; Figure 3); but the cooler the rearing temperature, the steeper the decline in locomotion toward high temperatures (Tukey posthoc tests indicates *β*_C 16°_ ≤ *β*_C 23°_ ≤ *β*_C 28°_ with *β*_C 16°_ < *β*_C 28°_; Figure 3; Supplementary Table S4). Despite considerable overlap in slope estimates among strains for the three rearing temperatures, especially at the hotter portion of the TPC, we also observe heritable strain differences in *β* in response to rearing conditions (Figure 3). For example, strain JU1338 exhibits virtually identical values of *β*_C_ regardless of the rearing temperature (similar for QX1547 and JU1348), whereas *β*_C_ values range 3-fold depending on rearing temperature for GXW23 (similar for HK104 and QR25). As we also described for *α*, these findings implicate heritable variation for phenotypic robustness of the dynamic behavioral response encapsulated by *β* as a function of rearing conditions. Even though both *α* and *β* derive from the same locomotory TPC, distinct genotypes can show robustness or plasticity separately for each trait, suggesting that these behavioral responses involve integration of the same environmental inputs with distinct genetic pathways.

The inflection point parameter (*τ*) identifies the temperature at which the TPC flips from concave to convex, heuristically indicating the center of the transition from normal to stopped locomotion. For all strains, low rearing temperatures led to colder values for the lower inflection point (*τ*_C_; ANOVA F_2,66_=215.75, P<0.0001) (Figure 3). However, the upper *τ* values (*τ*_H_) are similar regardless of whether worms were reared at 23°C or 28°C (*τ*_H_; ANOVA F_2,66_=39.29, P<0.0001, Tukey’s posthoc test indicates no significant difference between 23°C or 28°C), and 5 strains actually have a colder value of *τ*_H_ when reared at 28°C than 23°C (non-overlapping 95% CI for JU1338, HK104, EG4181, JU1348, NIC20). This suggests that limits to tolerance for high ambient temperature might not permit further upward shifts of the TPC with higher rearing temperature (Figure 3). Plasticity in *τ* as a function of rearing conditions is pervasive, with very few strains showing “robust” insensitivity to rearing temperatures (e.g. VT847 for *τ*_C_, JU1908 and VX34 for *τ*_H_; Figure 3). Temperate clade strains consistently show the greatest plasticity for *τ*_C_ in response to different rearing temperatures, with nearly a 4°C shift among rearing treatments, whereas the Nairobi strains exhibit consistently high plasticity for *τ*_H_ (Figure 3).

The TPCs from animals reared at cooler temperatures exhibit a ~1°C wider span between the two inflection points (Δ *τ*= *τ*_H_ - *τ*_C_), shifting position more on the cool end of the TPC and thus reflecting greater lability in response to cold ambient temperatures (ANOVA F_2,66_=6.39, P=0.0029, Tukey’s posthoc tests indicate Δ *τ*_16°_ = Δ *τ*_23°_ > Δ *τ*_28°_; Figure 3; Supplementary Table S4). Likewise, most strains have a narrower TPC (smaller Δ *τ*) when reared at 28°C than when reared under cooler conditions (Figure 4). The widest TPCs across the wild isolate strains are found consistently in those strains from the Temperate clade when reared at 16°C or 23°C (Figure 4), which could be interpreted as a feature of a “generalist genotype.” The TPC midpoints shift even more dramatically for the different rearing temperatures (ANOVA F_2,66_=157.95, P<0.0001, Tukey’s posthoc tests indicate significant differences among all rearing temperatures; Figure 4).

**Figure 4.**
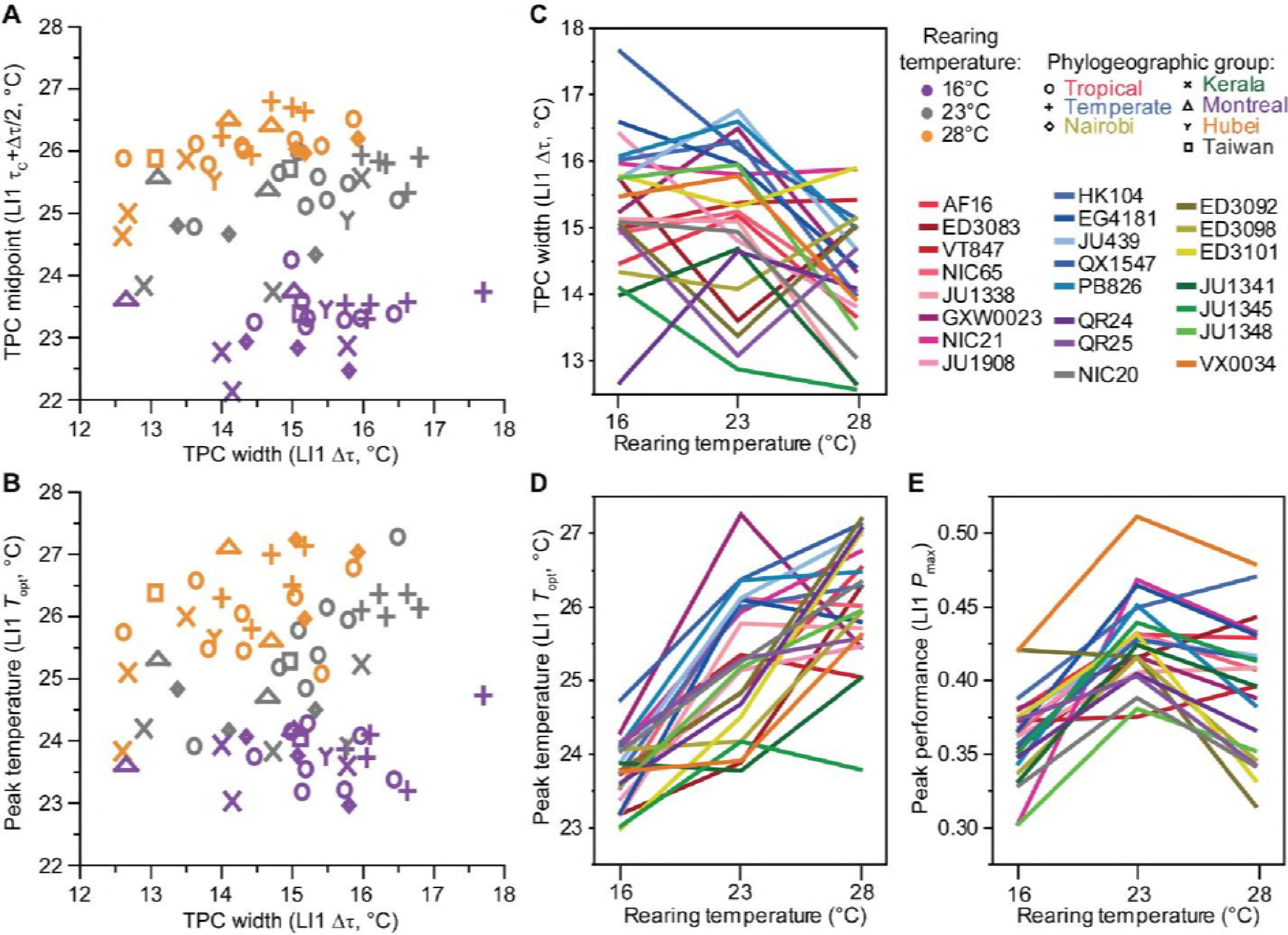
(**A**) *C. briggsae* thermal performance curve midpoint shifts warmer with higher rearing conditions (locomotion index LI1) (ANOVA F_2,66_=157.95, P<0.0001, all conditions differ by Tukey’s post-hoc test). The midpoint of the TPC also correlates positively with its width across wild isolate genotypes, for benign and hot rearing temperatures (Spearman’s ρ = 0.64, P=0.001 for 28°C; ρ = 0.47, P=0.024 for 23°C; ρ = 0.22, P=0.3 for 16°C). (**B**) The temperature of peak performance (*T*_opt_) correlates positively with TPC width across genotypes reared under benign or high temperature conditions (Spearman’s ρ=0.75, P<0.0001 for 23°C; ρ=0.41, P=0.051 for 28°C; ρ=-0.046, P=0.84 for 16°C). (**C**) TPC widths (Δτ) are narrower on average for wild isolate strains reared at 28°C than at lower temperatures (ANOVA F_2,66_=6.39, P=0.0029, Tukey’s post-hoc test). (**D**) The temperature of peak *T*_opt_ is greatest for wild isolate strains reared at high temperature (ANOVA F_2,66_=55.33, P<0.0001, Tukey’s posthoc tests: *T*_opt 28°_ > *T*_opt 23°_ > *T*_opt 16°_). (**E**) The peak performance for locomotory activity (LI1) *P*_max_, however, occurs for wild isolate strains reared under benign intermediate rearing conditions (ANOVA F_2,66_=20.25, P<0.0001, Tukey’s posthoc tests: *P*_max 23°_ > *P*_max 28°_ > *P*_max 16°_).

Much of the thermal performance curve literature emphasizes peak performance. Most strains reach their peak performance (*P*_max_) when reared at 23°C rather than the other two temperatures (ANOVA F_2,66_=20.25, P<0.0001, Tukey’s posthoc tests indicate *P*_max 23°_ > *P*_max 28°_ > *P*_max 16°_) (Figure 4); this is similar to the pattern for the asymptote parameter (*α*) with respect to rearing temperature. Just three strains have faster peak locomotion at 28°C instead of 23°C (ED3083, VT847, HK104; Supplementary Figure S3). To estimate the temperature for peak locomotion (*T*_opt_), which the logistic function fit does not capture, we fit a Gaussian curve (Equation 2) to locomotion data between 18°C and 32.5°C where the asymmetry in TPC shape is least pronounced (Figure 2). The temperature of peak performance (*T*_opt_) is warmest when we reared animals at 28°C compared to 16°C (ANOVA F_2,66_=55.33, P<0.0001, Tukey’s posthoc tests indicate *T*_opt 28°_ > *T*_opt 23°_ > *T*_opt 16°_), although for many strains there is little difference between rearing at 23°C versus 28°C (Figure 4). This is similar to the pattern we see with inflection point values (*τ*) and lends support to the idea that rearing temperature acts to shift the entire curve. We observed no significant correlation between peak performance *P*_max_ and the temperature at which peak performance occurred (*T*_opt_) (all P>0.53). We also found no significant correlation between peak behavioral activity (*P*_max_) and TPC breadth (Δ*τ*) across the 23 wild isolate genotypes for any rearing temperature, despite a positive trend for animals reared under benign 23°C conditions (Spearman’s ρ=0.38, P=0.073). By contrast, we observed significantly higher temperatures of peak performance (*T*_opt_) for genotypes that have broader temperature breadths (*τ*), provided the animals were not reared under cold conditions (Spearman’s ρ=0.75, P<0.0001 for 23°C rearing; ρ=0.41, P=0.051 for 28°C rearing; ρ=-0.046, P=0.84 for 16°C rearing).

### Robustness of thermal performance among individuals within a genotype

Our plots of TPCs exclude error bars to reduce visual clutter, but we also explicitly considered variation within strains and how this variation responds to assay temperature and to rearing temperature to learn more about individual thermal responses. Of course, the lowest standard deviation (SD) in locomotory activity among individuals occurs when all worms are paralyzed at the temperature extremes (Figure 2; Supplementary Figure S2). The SD is highest near temperatures that are slightly more benign than the inflection points in the TPC, where locomotion just begins to decline (Figure 2). However, high temperatures induce greater inter-individual variability for a given genotype as some worms stop moving entirely while others continue relatively normal locomotion, being most pronounced when we reared the animals at 28°C (Figure 2), which may associate with a wider range of individual body condition. The lower overall SD toward the cold end of the TPC indicates that the decline in locomotion tends to be more consistent across individuals, with this more stereotyped reaction indicative of a more canalized behavioral response to cooling than to warming ambient temperatures.

### Genetically distinct TPCs of Tropical and Temperate strains

We next sought to characterize differences between the “Tropical” AF16 and “Temperate” HK104 strains in detail, as the parental genotypes of a library of recombinant inbred lines with which we could explore the genetic architecture of trait variation. When reared at 23°C, the locomotory behavior of these two strains differed most at high temperature near 36°C, also reflected by the change in proportion of animals moving, despite similar locomotory activity of the two strains in benign and cool portions of the TPC (Figure 5; Supplementary Figure S4). The LI7 locomotion index metric has greater resolution than LI1 when animals are moving slowly (Stegeman *et al.* 2019), and LI7 reveals that near 36°C many individuals are moving slowly in AF16 whereas HK104 ceases locomotion altogether. This distinction in performance near 36°C is consistent across four large repeated experiments (Supplementary Figure S4). An instantaneous jump in temperature from a benign 23°C to 35.6°C also showed AF16 and HK104 to respond distinctively: the movement for AF16 worms declined more quickly over time, but retained slow activity for the duration of the ~21 min assay (Figure 5). By contrast, locomotion of HK104 declined more slowly toward complete immobility after ~13 min of exposure to 35.6°C (Figure 5), suggesting that AF16 can tolerate 35.6°C longer than HK104 can. Because of these consistent and conspicuous differences in high temperature responses for which the Tropical AF16 strain outperforms the Temperate HK104 strain, we devoted further effort to this aspect of high temperature locomotory performance.

**Figure 5.**
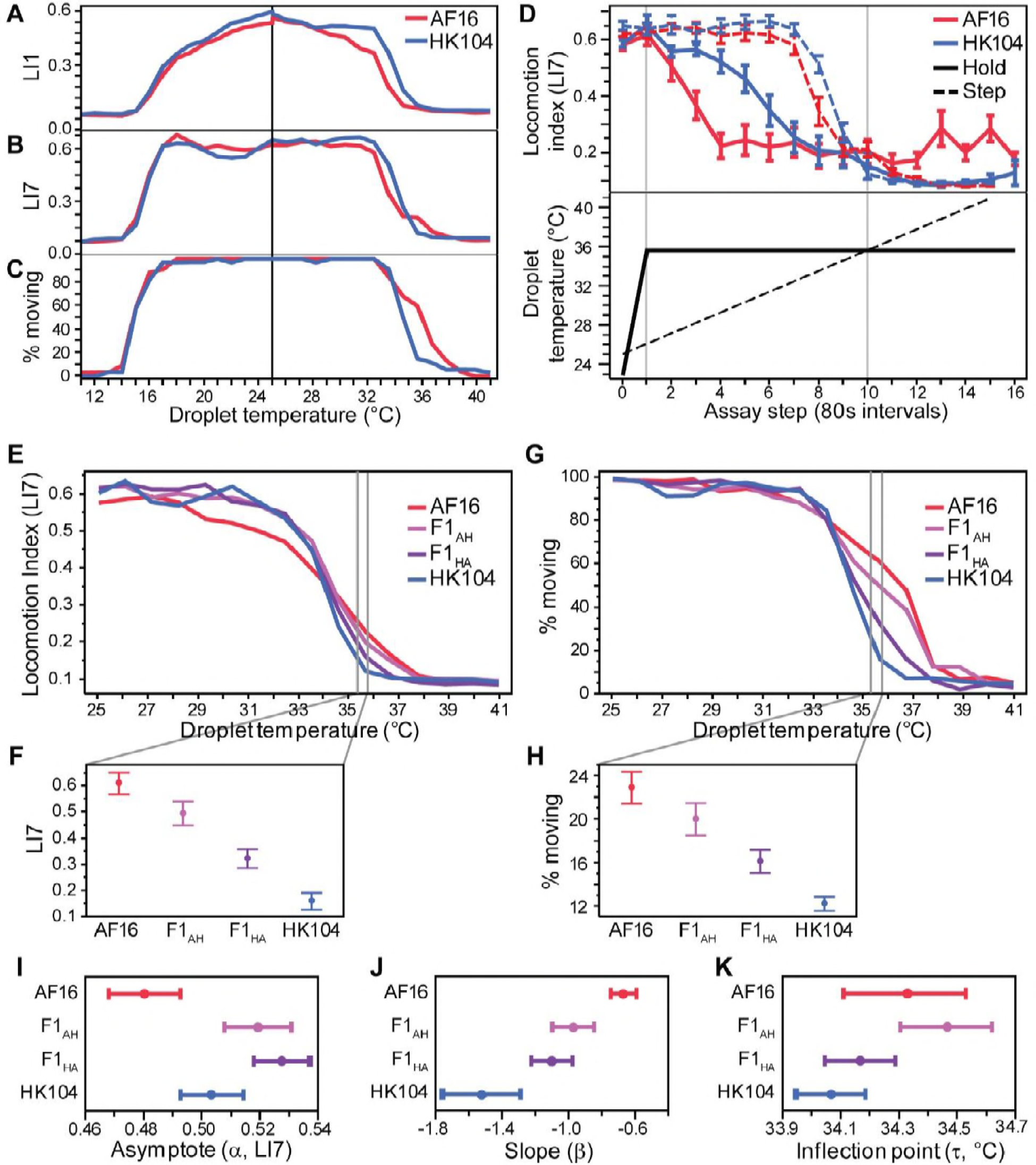
Thermal performance for Tropical strain AF16 (red) and Temperate strain HK104 (blue). (**A**) Decrementing and incrementing temperature TPCs show decline of locomotory performance with more extreme temperatures with three distinct metrics: (**A**) locomotion index with 1/15^th^ second resolution (LI1), (**B**) locomotion index with 7/15^th^ second resolution (LI7), and (**C**) the proportion of moving individuals. Upper and lower halves of the TPC start at an initial 25°C assay temperature (vertical black line). (**D**) Thermal responses for AF16 and HK104 with increasing temperature steps (dashed lines, same data as in A) or with an instantaneous increase from 23°C to 35.6°C followed by a constant hold temperature (solid lines); top panel shows TPCs and bottom panel shows experimental thermal profiles). Error bars indicate ±1SEM. (**E**) Comparison of locomotion index (LI7) TPC and (**F**) values at 35.6°C (±1SEM) for F1 heterozygotes to their homozygous parental genotypes (Tropical AF16 and Temperate HK104; purple = F1_HA_ with HK104 as maternal parent; pink = F1_AH_ with AF16 as maternal parent). (**G**) Comparison of proportion moving TPC for F1 and parental genotypes, with (H) point estimates at 35.6°C (±1SEM). (**I-K**) Parameter estimates from logistic function fits to LI7 TPCs for F1 heterozygotes and parental genotypes show more extreme values for baseline locomotion (α) in heterozygotes and intermediate F1 values for thermal sensitivity (β). All animals reared at 23°C.

The reciprocal F1 progeny of AF16 and HK104 exhibited largely intermediate TPC phenotypes at those high temperatures that most distinguish the parental strains (35.6-36.7°C), with the F1s appearing slightly more similar to their maternal parent to indicate some parent-of-origin asymmetry (Figure 5). These F1 data cannot distinguish mitochondrial versus epigenetic parental effects, although we can exclude sex-linkage as a factor because all individuals were hermaphrodites (XX karyotype); QTL mapping with RILs found no evidence for a mitochondrial contribution (see below). The largely intermediate F1 heterozygote phenotypes imply that trait variation cannot be explained by a single dominant gene of large effect, although the inflection point parameter values are more ambiguous on this point (Figure 5). These results also tell us that heterozygous genotypes do not exhibit outbreeding depression for locomotion in response to increasing temperature, in contrast to some other phenotypes in *C. briggsae* (Dolgin *et al.* 2007), with the asymptote parameter actually showing overdominant trait values that exceed both parents (Figure 5). We next applied quantitative trait locus mapping using recombinant inbred lines to further dissect the genetic architecture of these TPC differences.

### Locomotory thermal performance variation among recombinant genotypes for QTL mapping

We quantified locomotory thermal performance for 153 RILs, plus their two parental lines, with the liquid microdroplet assay for the ‘hot’ portion of the thermal performance curve. We then tested for associations between genotype and phenotype, yielding multiple QTL across the genome that each explain up to *H*^2^ = 27.6% of the variation in behavioral thermal performance traits (Supplementary Table S5). We summarized trait variation in temperature-dependent locomotory responses using a diversity of phenotypic metrics and reduced their dimensionality through function-valued trait analysis of the reaction norms and Principal Component Analysis (PCA). Most RILs produced phenotypes that spanned the trait space between parental values, but those that fell beyond parental performance indicate transgressive segregation and imply multiple loci controlling trait variation. For example, 52 RIL strains have steeper point estimates for slope (|*β*_H_|) than the HK104 parent whereas only 7 RILs had a shallower estimated slope than AF16 (Figure 6). Similarly, only 6 RILs showed higher locomotory activity than AF16 at 35.6°C (LI7) and only 9 RILs have inflection point values (*τ*_H_) higher than AF16 (Figure 6); *β*_H_ and *τ*_H_ correlated only weakly across RIL strains (*r*=0.127 *P*=0.0132), suggesting that the genetic basis to variation in these traits may be distinct. Many strains (83) have asymptote estimates lower than either parent, indicating that they have slower baseline locomotion (*α*_H_; Figure 6). Overall, the range of variation among the RILs is similar to (or greater than) that seen among wild isolate genotypes of *C. briggsae*: mixing up the genomes of just two genotypes to create the RILs produced a range of phenotypic variation similar to the natural range of variation seen in the entire species (Figure 6).

**Figure 6.**
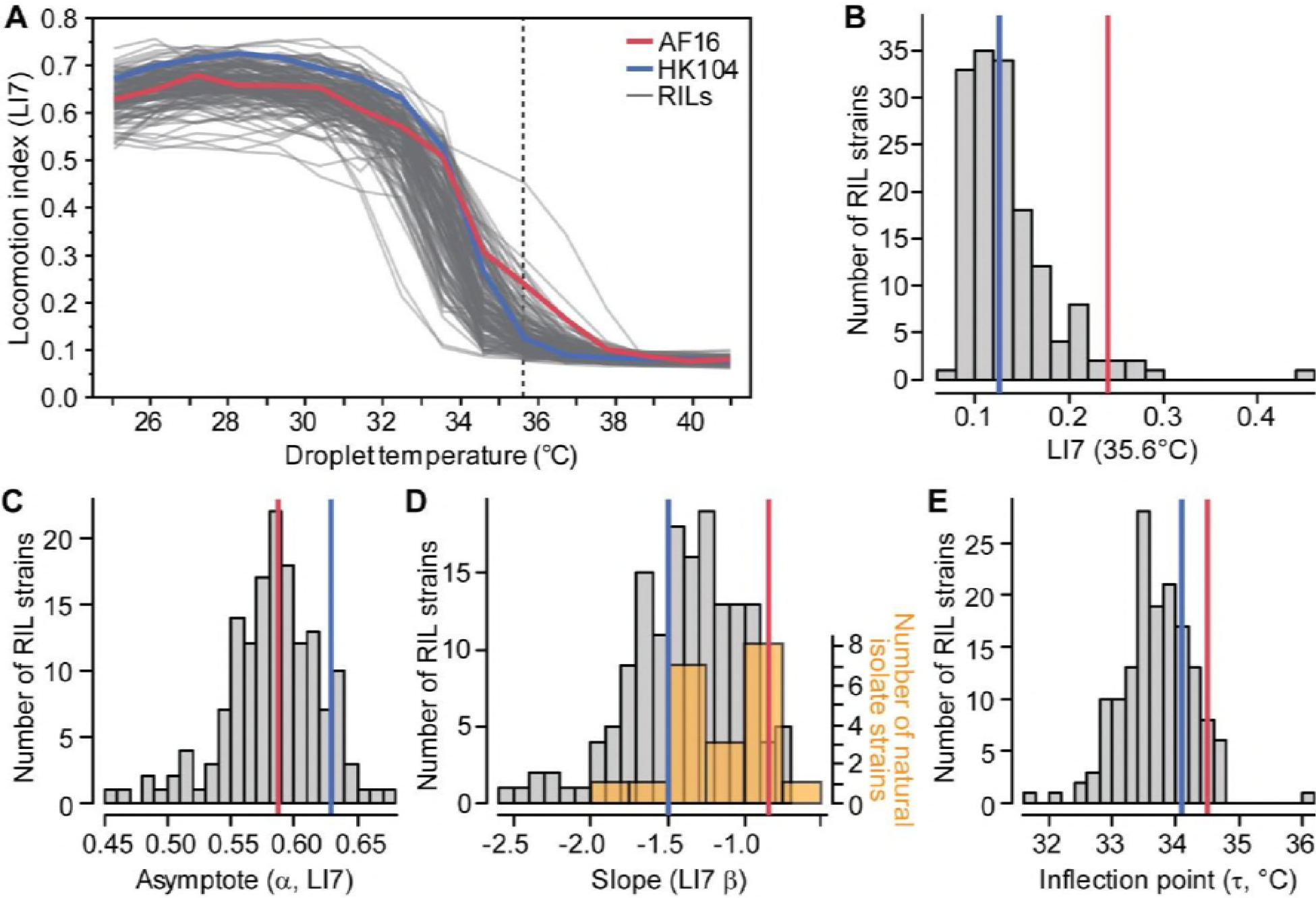
(**A**) Thermal performance curves for 153 recombinant inbred lines (RILs) (gray) derived from *C. briggsae* parental genotypes (Tropical AF16, red; Temperate HK104, blue). Locomotion indices (LI7) for each strain at 35.6°C (dashed vertical line) is shown as a histogram of phenotypic values in (**B**). (**C-E**) Distribution of parameter estimates from function fits of locomotion index scores to each strain (LI7; see Equation 1 and Figure 1). Distribution of slope estimates (β_H_) for 23 wild isolate strains (orange) in (D) is superimposed on the distribution for the 153 RILs (gray) for function fits to the LI7 metric from TPCs of worms reared at 23°C. Red and blue lines show values for strains AF16 and HK104.

A function-valued trait approach provides one powerful way to distill a dynamic multivariate phenotype down to a simpler representation (Stinchcombe and Kirkpatrick 2012). We also applied PCA to reduce the dimensionality of the 167 metrics for the 155 strains to yield 8 principal components that are statistically independent and together explain two-thirds of the total variation (Supplementary Figure S5). Correlated phenotypes get weighted strongly together in a given principal component (PC) axis to represent an integrated “synthetic” phenotype, and we analyzed this subset of 8 PCs as composite traits for subsequent analysis. The first principal component (PC1) explains ~20% of the phenotypic variability and relates primarily to locomotion at low to moderate temperatures up to 32.5°C, where locomotion is normal or just starts to decline for many strains (Figure 7; Supplementary Figure S6). Subsequent orthogonal components explain smaller portions of phenotypic variation and become more dominated by a few phenotypes with high weights (Supplementary Figure S6). The variation in PC2 captures ~16% of the variation among the RIL phenotypes, generally being associated with responses to high temperatures above 35.6°C (Figure 7; Figure 8; Supplementary Figure S6). We also included measures of within-strain variation itself as “penetrance” phenotypes: the standard deviations for locomotion metrics and standard error for parameter estimates. These penetrance metrics tend to cluster together in PCs 3, 4, 6 and 7 that are distinct from the PCs loaded primarily with simple locomotion metrics (PCs 1, 2, 5; Supplementary Figure S6).

**Figure 7.**
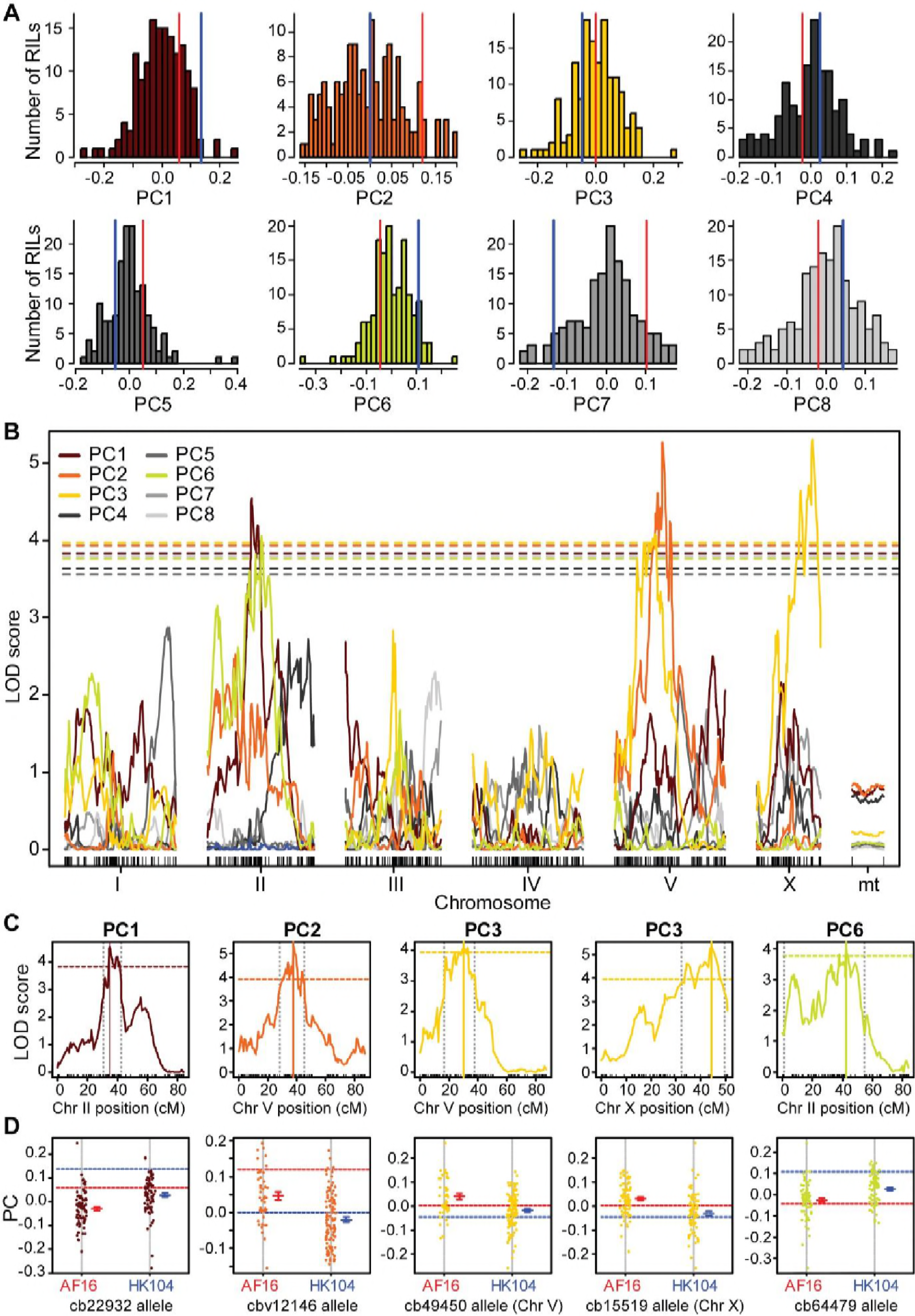
(**A**) Distributions of ‘synthetic’ locomotion behavior trait values for 153 RIL strains for the first eight principal components (PCs). Red lines indicate AF16, blue lines indicate HK104. (**B**) QTL maps for the 8 ‘synthetic’ principal component locomotion behavior traits show logarithm of the odds (LOD) scores plotted against marker position by chromosome (“mt” = mitochondrial genome). Dashed lines indicate the 5% genome wide significance threshold from 2500 permutations and Bonferroni correction (P=0.05/8). PCs with non-significant QTL indicated with gray colors in both (A) and (B). (**C**) Significant QTL peaks shown for individual chromosomes (colored as for PC traits shown in B; significance thresholds as in B). Solid vertical lines indicates LOD peak for the marker loci shown in (D); dashed vertical lines indicate the 95% Bayes credible interval. (**D**) Phenotype values of RIL strains segregated by parental genotype at the marker closest to the QTL peak in (C) indicated above each panel. Parental strain values indicated by horizontal dashed lines (red AF16, blue HK104).

**Figure 8.**
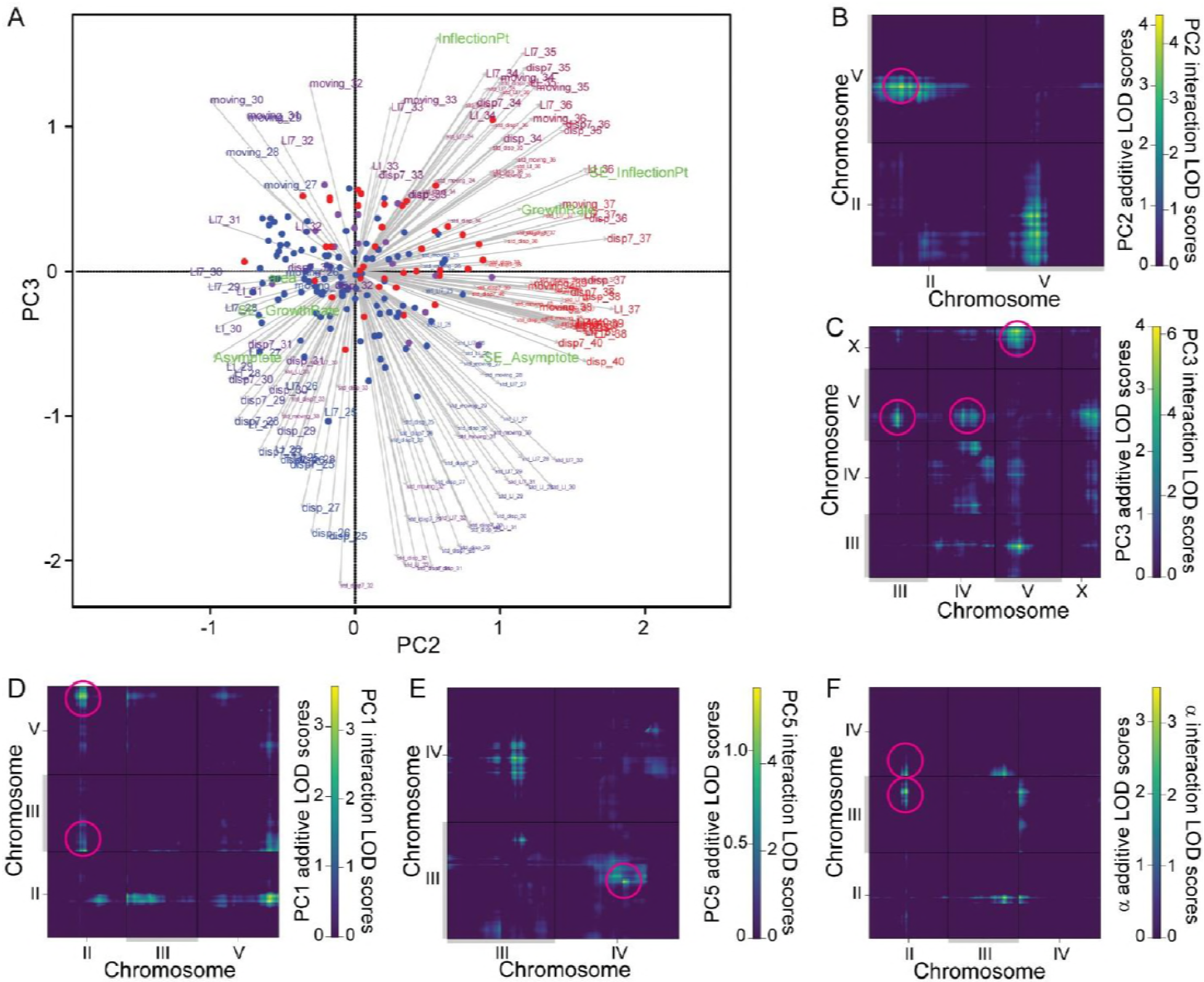
(**A**) Biplot of 167 individual phenotype weights measured for the 153 RILs, for the orthogonal PC2 and PC3 ‘synthetic’ trait axes that map to Chromosome V. Length and direction of vectors indicate weight of the phenotype in each PC with the text indicating individual phenotype metrics (Supplementary Figure S6). For example, the Locomotion index (LI7) at 35.6°C is weighted strongly in both these PC’s and can be found near the top-right corner (“LI7_35”). Circles indicate each RIL strain, colored according to their genotype for markers at QTL peaks for PC2 and PC3 on Chromosome V (marker cbv12146 for PC2 and marker cb49450 for PC3; red = both AF16 alleles, blue = both HK104 alleles, purple indicates different alleles between the peak of PC2 and PC3; peaks were 6cM and ~3.9Mb apart, Supplementary Table S5). (**B-F**) Two-dimensional QTL scans show pairwise LOD scores for a given phenotype, shown only for chromosomes with significant QTL. The upper-left triangle above the 1:1 line in each plot contains the additive-effect LOD values after subtracting the LOD values from the single-QTL model. The lower-right triangle below the 1:1 line in each plot shows the LOD values for interactions. Note different color scales in each plot and each off-diagonal triangle zone. Magenta circles highlight the significant QTL locations (Supplementary Table S5).

To perform QTL mapping, we integrated the RIL phenotype data with the RIL genotypes for 1031 SNP markers (Ross *et al.* 2011), which comprised 430 distinct genetic blocks in the genomes of the 153 RILs used here. We first mapped the eight principal component axes as “synthetic traits”, using 2500 permutations to determine genome-wide significance thresholds based on a Bonferroni correction for eight phenotypes. This reduction in data dimensionality allowed the maintenance of power to detect QTL while avoiding excessive multiple hypothesis testing and overly redundant phenotypes among the 167 metrics of temperature-dependent locomotory behavior. We then validated the biological interpretations of the “PC-QTL” with genomic associations involving a key univariate trait (LI7 at 35.6°C). However, we focus our QTL mapping analysis and interpretation on the three TPC function fit parameters for the ‘hot’ portion of the thermal performance curve rather than PCs (slope *β*_H_, asymptote *α*_H_, inflection point *τ*_H_). These analyses uncovered at least one QTL on chromosome V that explains *H*^2^ =12-28% of the variation in locomotion at high temperature (depending on the specific phenotype) and another major QTL on chromosome II associated with locomotion at lower temperature (*H*^2^ = 12-19%; Supplementary Table S5). The function fit parameters as phenotypes are especially interesting because they give a summary of characteristics of the entire thermal performance curve and correspond strongly to some of the less intuitive summaries of phenotype captured with PCs.

### Locomotion at benign temperature maps to QTL on Chromosome II

Four of the eight synthetic PC traits mapped significantly to QTL (Figure 7). Specifically, both of the synthetic traits corresponding to locomotion at benign initial temperatures in the liquid micro-droplet assay, PC1 and PC6, as well as the asymptote parameter (*α*_H_) map to the center of Chromosome II (Figure 7; Figure 9). Other univariate phenotypes for locomotion at benign temperatures also had LOD peaks on Chromosome II in exploratory univariate analyses (Supplementary Figure S7; Supplementary Figure S8), many of which weigh heavily in PC1 and/or PC6 (Supplementary Figure S6). These QTL on Chromosome II may have a common genetic basis that affects all of these phenotypes. The HK104 allele at the marker nearest to the QTL peak is correlated with higher values of locomotion metrics (Figure 7; Figure 9), consistent with the parental strain differences (Figure 5).

**Figure 9.**
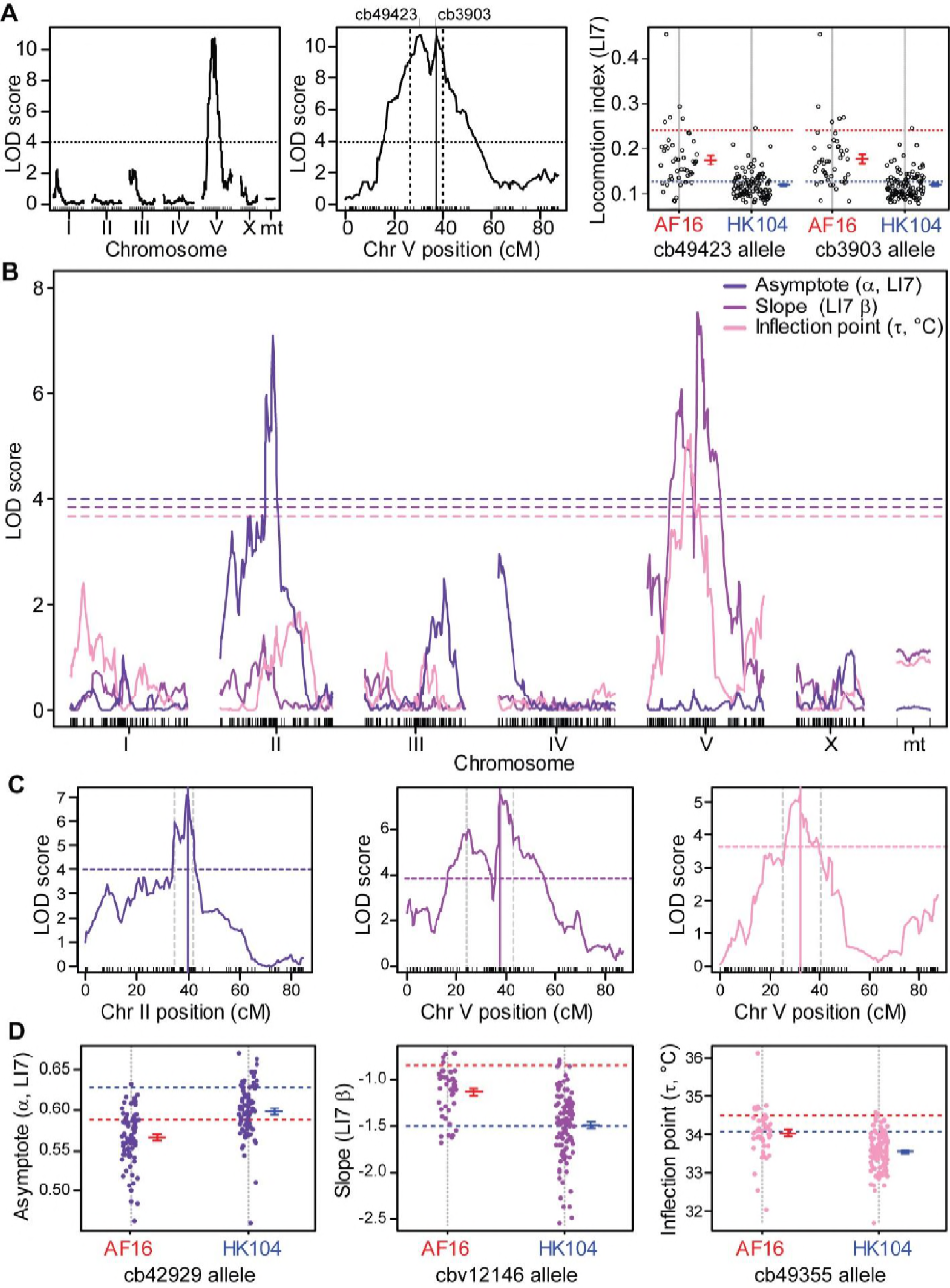
(**A**) Multiple imputation QTL mapping analysis of the Locomotion index (LI7) phenotype at 35.6°C, for all chromosomes and an expanded plot for the QTL on Chromosome V. LOD peak on Chromosome V at 37.35cM indicated by solid vertical line, with dashed vertical lines indicating 95% Bayes credible interval. Horizontal lines indicates the 5% genome-wide significance LOD threshold after 2500 permutations and Bonferroni correction as in Figure 7. Panel on right shows phenotype values of RIL strains segregated by parental genotype at the markers closest to the QTL peaks on Chromosome V (mean ±1SE for each marker allele shown adjacent to the distribution of points); values for parental strains shown as dashed horizontal lines (red AF16, blue HK104). (**B**) QTL mapping for three function fit parameter phenotypes for locomotion index (normalized LI7). Dashed lines indicate threshold for significance from 2500 permutations of the data (Bonferroni corrected P=0.05/12). (**C**) LOD peaks (solid lines) and 95% Bayes credible intervals (vertical dashed lines) for chromosomes with significant QTL for each function parameter phenotype; significance thresholds as in (B). (**D**) Phenotype values of RIL strains segregated by parental genotype at the marker closest to the QTL peak in (C) indicated above each panel. Mean ±1SE across RILs for each marker allele shown adjacent to the distribution of points; dashed horizontal lines indicate parental strain phenotypes (red AF16, blue HK104). See Supplementary Table S5 for physical map positions of QTL.

### High temperature locomotory activity maps to QTL on Chromosome V

Two synthetic PC traits mapped as QTL to the center of Chromosome V (PC2 and PC3), and PC3 also mapped to a QTL on the right end of the X Chromosome (Figure 7). The QTL peaks for PC2 and PC3 on Chromosome V are 3.9Mb apart, with partially-overlapping 95% Bayes credible intervals. Univariate phenotypes related to locomotion at high temperatures (>35°C) load most heavily on PC2, whereas PC3 is most closely tied to “penetrance” phenotypes related to variability among individuals at moderately high temperatures (<33°C; Figure 8, Supplementary Figure S6). Consequently, we mapped the univariate 35.6°C locomotion index (LI7), also yielding a QTL peak on Chromosome V, in fact the highest LOD value for any phenotype (Supplementary Table S5), and consistent with the biological interpretation of at least one QTL on Chromosome V associated with high temperature locomotory activity. Additional exploratory univariate QTL analyses reinforce this interpretation, given that 32 univariate traits related to high temperature locomotion associate with this region of Chromosome V even after Bonferroni multiple-test correction (Supplementary Figure S6; Supplementary Figure S9). Moreover, the function-valued trait parameters for slope (*β*_H_) and inflection point (*τ*_H_) also both mapped to QTL in the center of Chromosome V (Figure 9), further indicating that this genomic region contains important genetic variation explaining locomotory differences at high temperatures. Strains with Tropical AF16 genotypes at the peak locations of these QTL had values related to greater performance at high temperature, consistent with the difference between parental strains (Figure 7; Figure 9).

The QTL on the X chromosome for PC3, and exploratory analysis with univariate metrics, mostly corresponds to phenotypes related to within-strain standard deviation for locomotion at relatively benign temperatures (Supplementary Figure S6; Supplementary Figure S7). One possibility is that genetic variation on the X chromosome contributes to the robustness of locomotory responses among individuals following exposure to moderately increasing temperatures.

Given the important role implicated by the Chromosome V QTL for behavioral performance at high temperatures, we sought to affirm a causative role for genetic differences between Tropical AF16 and Temperate HK104 in shaping locomotory thermal performance. Therefore, we created 13 NILs that had regions of HK104’s Chromosome V introgressed into the genomic background of AF16 by backcrossing RILs to AF16 for 7-12 generations using marker-assisted selection. We observed significant differences in locomotion among the NILs at 35.6°C, most of which exhibited phenotypes more similar to the HK104 parent (Supplementary Figure S10). Consistent with QTL on Chromosome V, therefore, it can be sufficient for HK104 genotypes in the center of Chromosome V of the NILs to shift the phenotype toward HK104-like trait values (e.g. A1124-gs123 and A1124-gs124). Despite failing to narrow the QTL region, these introgression lines affirm the QTL region on Chromosome V as containing genetic differences that cause phenotypic differences in behavioral thermal performance at high temperatures.

The QTL region for β and τ spans nearly 11Mb and contains 2837 putative genes; the Bayes interval for locomotion (LI7) at 35.6°C spans almost 7Mb and contains 1784 putative genes. Approximately 70% of these genes have associated gene ontology (GO) terms, implicating potential structural or molecular functions that we cross-referenced with genes for which we detected coding changes (missense, nonsense, or indel alleles). We identified 115 of the 2837 genes (63 of the 1784 genes) to have such coding changes as fixed differences between 20 Tropical and 4 Temperate group strain genomes (Thomas *et al.* 2015) (Supplementary Figure S11, Supplementary Table S7). Fixed differences between these phylogeographic strain groups should represent changes that are most ecologically relevant to temperature-dependent phenotypic divergence. Sensory and neurological system genes, including membrane proteins, receptors signaling pathway genes were significantly overrepresented relative to *C. briggsae*- wide expectations (PANTHER overrepresentation test; Supplementary Figure S11) (Thomas *et al.* 2003). Despite the enrichment for these genes with potential influence on behavior, many other kinds of genes could conceivably contribute, in addition to the potential influence of non-coding regulatory or RNA genes on thermal performance.

### Testing for multiple QTL on Chromosome V

The LOD peaks on Chromosome V visually appear bimodal for *β*_H_ and LI7 at 35.6°C, and the broad Bayes credible interval suggests that multiple causal loci might occur under the QTL region (Figure 9). The primary peak is located between 37.4 and 38.0 cM (near the QTL peak for PC3) and the secondary peak across the “valley” is between 25.0 and 31.5cM (near the QTL peak for PC2) (Supplementary Figure S9). Presuming that the two PCs capture distinct orthogonal aspects of the phenotypes, then the ‘twin peak’ pattern on Chromosome V might hint at distinct genetic contributions to temperature-sensitive traits. Can we find statistical evidence for twin peaks? It was not possible to map them separately with the multiple imputation method used for our primary analysis, which assumes only one peak per chromosome, prompting us to consider alternate approaches. Only 15 of the 153 RILs have genotypes that differ between the 37.4cM and 31.5cM locations, however, suggesting limited power to disentangle separate loci. Nevertheless, we explored the possibility of twin peaks using two-dimensional and multidimensional QTL analyses, which did not reveal evidence of a significant pair of QTL peaks on Chromosome V (Supplementary Figure S12).

As an alternate approach, we performed two-dimensional QTL scans, which found evidence for possible new additive and interacting QTL that the single-QTL model could not detect (Figure 8; Supplementary Table S6). These two-dimensional scans test all pairwise combinations of intervals at least 5cM apart for interactions and epistatic effects for 10 key phenotypes, including those with significant QTL from our initial analysis (those in Supplementary Table S5). However, this procedure did not reveal any new QTL for the phenotypes PC6, slope (β), inflection point (τ) or locomotion (LI7) at 35.6°C. Instead, the two-dimensional QTL analysis revealed only one pair of loci with an epistatic interaction and eight pairs of loci with additive effects across different chromosomes for the remaining phenotypes (Figure 8; Supplementary Table S6). For PC1 and asymptote (α), this analysis recapitulates QTL at the center of Chromosome II. At least three QTL were also found on Chromosome V associated with the ‘synthetic’ phenotypes PC1, PC2 and PC3. Interestingly, PC2 shows evidence for an additive interaction between QTL near the centers of Chromosomes II and V, which correspond to the two distinct QTL from the single-QTL analysis for the two separate phenotypes.

## Discussion

### Behavioral phenotypic plasticity in response to temperature

Phenotypic reaction norms inherently illustrate the plasticity of traits in response to extrinsic conditions (Via *et al.* 1995; Dingemanse *et al.* 2010). Our analysis of locomotory behavior in response to thermal inputs show this phenotypic plasticity explicitly, reflecting both dynamic changes in temperature over time and the plastic responses of those dynamic reaction norms to distinct rearing conditions. We summarized this complex kind of phenotype with a function-valued trait approach (Wu and Lin 2006; Stinchcombe and Kirkpatrick 2012), using biologically interpretable parameter values from fits of a logistic function to the norms of reaction. This strategy distilled a highly multidimensional phenotype into three key components captured by: the baseline locomotory activity of animals at benign temperatures (asymptote parameter α), the rate of change in motility with changing temperature (slope parameter β), and the temperature mid-way through the transition from normal movement to immobility (inflection point parameter τ).

Our characterization of behavioral dynamics with changes in temperature revealed *C. briggsae* locomotory behavior to produce a response similar to a classic thermal performance curve (TPC) (Huey and Stevenson 1979; Huey and Kingsolver 1989). The classic TPC shape of a steeper decline in trait performance toward hot temperatures from a peak at benign temperatures, relative to the shallower trait change toward cool temperatures, was most pronounced when animals were reared under cool conditions. Such cool rearing conditions also produced TPCs with a broader range of animal activity (larger Δτ), indicative of a more ‘generalist’ trait response to changes in the ambient thermal regime. Thus, ‘developmental acclimation’ (Colinet and Hoffmann 2012) from early life experience cascades into producing adult animals with more generalist or specialist locomotory strategies. The greater breadth of the behavioral response profile, however, primarily reflected expansion of locomotory activity when experiencing cool ambient temperatures, both in terms of a cold-shifted inflection point (τ) and a shallower slope (|β|). By comparison, the upper ‘hot’ portion of the TPC was much less sensitive to the rearing conditions of animals, indicating that the temperature regime that worms experience early in life is disproportionately important in shaping their response to cooling ambient temperatures. This finding of greater TPC plasticity in a cool regime is generally consistent with studies of thermal biology in other systems (Chown and Terblanche 2006). Previous work in *C. elegans* demonstrated that nematode TPC responses to high temperatures are influenced, in part, by neural ‘decision making’ and that the sensitivity of TPCs to rearing conditions also has a genetic component (Stegeman *et al.* 2019). This sensitivity of TPC shape to rearing conditions in both *C. elegans* and *C. briggsae* illustrates the profound influence that early life experience through developmental acclimation (Colinet and Hoffmann 2012) can exert on subsequent traits later in life, as is well-known in more complex animals like insects and humans (Urquhart-Cronish and Sokolowski 2014).

Despite the variation in TPC breadth, both due to rearing conditions and to natural differences in genotype, we found no evidence supporting the ‘jack of all temperatures and master of none’ hypothesis for the existence of specialist-generalist trade-offs due to functional constraints in thermal adaptation (Huey and Hertz, 1984). Specifically, we observed that TPC breadth (Δτ) and peak behavioral activity (*P*_max_) did not correlate negatively. Curiously, we also failed to find support for the ‘hotter is better’ hypothesis, an idea motivated by the logic that adapting to hotter environments would lead to a higher peak performance (Huey and Kingsolver 1989; Angilletta *et al.* 2010): *C. briggsae* genotypes with hotter temperatures of peak performance (*T*_opt_) did not actually show higher peak performance (*P*_max_). Instead, we find higher temperatures of peak performance (*T*_opt_) for genotypes that have broader temperature breadths (Δτ), which suggests that ‘generalist genotypes’ are more likely to have peak performance at high temperatures. The behavioral nature of our phenotypic analysis provides one interesting possible explanation for the absence of evidence to support these classic hypotheses from thermal biology. Specifically, because locomotory activity is controlled in part by neural decision-making (Stegeman *et al.* 2019), the optimal (adaptive) performance at a given temperature might not correspond to peak activity. The behavioral decision to slow movement as the animal senses more extreme temperatures might be the feature itself that is subject to selection, potentially in an adaptive manner.

Genetic mapping studies of plasticity have revealed gene-by-environment interactions for life history traits in diverse organisms, including *C. briggsae*’s nematode cousin *C. elegans* (Vieira *et al.* 2000; Ungerer *et al.* 2003; Hausmann *et al.* 2005; Gutteling *et al.* 2007). However, analysis of *Drosophila serrata* locomotory activity in thermal performance curves from natural variation (Latimer *et al.* 2011), QTL mapping with RILs (Latimer *et al.* 2015), and via mutation accumulation (Latimer *et al.* 2014) provides an especially useful comparison to our work on *C. briggsae*. Like our analysis for *C. briggsae*, Latimer and colleagues’ data did not support the “hotter is better” hypothesis, instead suggesting that tropical genotypes of *D. serrata* were more specialized, with a higher peak performance and narrower TPC than other natural populations. Most variation in TPCs among *D. serrata* RILs reflected overall locomotion, rather than changes in TPC shape or temperature-specific responses. By contrast, we observed for *C. briggsae* that all key characteristics of TPC shape that we analyzed differed significantly among genotypes.

In addition to the influence of temperature changes on the average response of animal movement, we also observed consistent patterns in worm-to-worm variability as a dynamic response to temperature for a given genotype. Cooler ambient temperatures generally led to more stereotyped behavior, consistent with the locomotory response being more canalized or robust to perturbations in temperatures below benign conditions. By contrast, worm-to-worm variation was greatest at hot temperatures just preceding the rapid decline in swimming behavior in the TPC, indicating greater individual stochasticity in behavioral responses as ambient heating conditions become more extreme. This pattern suggests that selection might have played a greater role in shaping plastic responses to cooling temperature in combination with the greater sensitivity of cool-regime TPCs to developmental rearing conditions, analogous to insect systems (Chown and Terblanche 2006).

### Heritable contributions to behavioral thermal reaction norms

By quantifying the thermal performance curves for 23 wild isolate genetic backgrounds of *C. briggsae*, we demonstrated substantial heritable variation in behavioral reaction norms. Essentially all of the features of locomotory responses to changing temperatures exhibited heritable differences among distinct genotypes from across the range of this species’ genetic diversity, from baseline or maximal activity (α; *P*_max_) to the sensitivity to temperature changes toward hot or cold (β_H_, β_C_), the temperature of maximum thermal sensitivity or performance (τ_H_, τ_C_; *T*_opt_), and thermal breadth of activity (Δτ). We then interrogated the ‘hot’ portion of *C. briggsae* behavioral thermal responses in more detail for representatives of the ‘Tropical’ and ‘Temperate’ phylogeographic groups within this species (Cutter *et al.* 2006; Felix *et al.* 2013; Thomas *et al.* 2015), followed by a library of 153 recombinant inbred lines (RILs) derived from these Tropical and Temperate parental strains (Ross *et al.* 2011). These latitudinally-separated genetic groups within *C. briggsae* show evidence of adaptation to temperature differences in terms of fecundity (Prasad *et al.* 2011), and also exhibit distinctive behavioral responses in terms of thermal preference and isothermal tracking (Stegeman *et al.* 2013).

The Tropical (AF16) and Temperate (HK104) genotypes showed distinctive responses to high temperature, in particular, with animals with the Tropical genotype continuing to move slowly above 35°C whereas Temperate strain worms were quiescent. Consistent with multiple loci contributing to this difference, we found that F1 individuals exhibited intermediate behavioral responses and that the recombinant genotypes of the RIL strains produced a spectrum of responses that exceeded the parental trait values, in fact yielding a similar range of phenotypic variation as we observed across the genetically diverse set of wild isolates. Our quantitative trait locus (QTL) mapping analysis corroborated this idea of multiple contributing loci, with major QTL mapped to three chromosomes.

Distinct components of the behavioral response to temperature correlate with genetic differences in unlinked portions of the genome. The baseline behavioral activity associates with the center of chromosome II (LOD peak at position 9.91 Mb for α_H_), whereas the behavioral sensitivities to high temperature map to chromosome V (11.62 Mb for β_H_, 8.08 Mb for τ_H_). We observed similar genetic architecture for alternate ways of summarizing phenotypes for the behavioral reaction norm, whether using univariate metrics or Principle Component axes as synthetic phenotypes, which corroborates the biological importance of loci on chromosomes II and V. Moreover, univariate and synthetic “PC-QTL” indicate that genetic variation on the X chromosome contributes to inter-individual variation in behavioral responses as temperature increases, affecting the robustness or canalization of the trait when perturbed by environmental changes to thermal regime. These genetically-separable components of the behavioral thermal performance indicate that it is encoded by distinct modules that together contribute to the overall complex behavioral response of the reaction norm. Independent genetic loci contributing to complex phenotypes in a modular way might be a general feature of behavioral phenotypes, as it also has been observed in tunnel burrowing behavior in oldfield mice (Weber *et al.* 2013).

Genetic perturbations are more likely to affect locomotion detrimentally, especially at high temperature, than they are to enhance locomotion or tolerance to extreme higher temperatures (Stegeman *et al.* 2019). Many genes must work in concert for successful locomotion, so even slight genetic changes to the system could decrease tolerance to extreme temperature stress and consequently result in slowed locomotion. With phylogeographic differentiation, co-adaptation may have tuned gene variants to work best in combination within their ‘home’ genome, especially in a self-fertile species like *C. briggsae* (Dolgin *et al.* 2007). By recombining genomes from genetically distinct phylogeographic groups, these co-adapted genetic networks would get disrupted. We see evidence suggestive of this idea here in that a preponderance of RILs slowed their locomotion in response to temperature shifts relative to both parental strains (Figure 6).

Despite the complexity of a dynamic trait like a behavioral response to external inputs, we uncovered little evidence of non-additive interactions and just a handful of QTL across the genome. This finding in *C. briggsae* contrasts with an epistatic genetic architecture to thermal preference behavior in *C. elegans* QTL analyses (Gaertner *et al.* 2012). We also found no contribution of mitochondrial differences, despite previous work in *C. briggsae* showing an influence of mitochondrial genotype on other features of their biology (Howe and Denver 2008; Howe *et al.* 2010; Ross *et al.* 2011; Chang *et al.* 2015). This lack of a role for mitochondria in our analysis also contrasts with mitochondria contributing to variation in thermotolerance in insects and rodents, owing to the possible contribution of mitochondrial membrane potentials releasing heat (Fontanillas *et al.* 2005; Ballard *et al.* 2007). Evidence of paternal inheritance of mitochondria in *C. briggsae*, especially those involving HK104 (Ross *et al.* 2016), might also mask our ability to infer mitochondrial QTL. Our QTL mapping analysis also suggests several additional potential QTL at the cusp of statistical significance that might contribute minor phenotypic effects to explain residual heritable variation that is not captured in the major QTL we identified on Chromosomes II, V and X. While our experiments with introgression line genotypes affirmed a causative role of the Chromosome V QTL in affecting behavioral thermal responses, fine-mapping the causative underlying genetic differences in this gene dense region with low recombination will require further enquiry (Stein *et al.* 2003; Ross *et al.* 2011; Thomas *et al.* 2015).

Given the distinct phenotypes that map to overlapping locations on Chromosome V and the hundreds of genes with potentially function-affecting differences between the Temperate and Tropical strains, it remains plausible that multiple linked loci within the QTL region contribute to the locomotory reaction norm. Variation in both β_H_ and τ_H_ mapped to QTL on Chromosome V, but these function parameter phenotypes are mostly independent of one another (*R*^*2*^=0.04, *P*=0.0132,), and the orthogonal PC2 and PC3 synthetic phenotypes also both map to Chromosome V. Moreover, dissociating multiple factors from one another would weaken the individual signal of each factor so as to make it challenging to decipher the role of the separated genetic loci. For example, the characterization of natural variation in *C. elegans*’ *srx-43* following QTL mapping of pheromone-dependent foraging follows this pattern of a more modest phenotypic influence of an isolated causal genetic factor (Greene *et al.* 2016). QTL mapping in *C. elegans* has identified multiple causal variants of large effect (Seidel *et al.* 2008; Ghosh *et al.* 2012), often associated with domestication to lab conditions (e.g. *npr-1*, *glb-5*, *nath-10*) or with a major functional disruption (e.g. *plg-1*) (Palopoli *et al.* 2008; Mcgrath *et al.* 2009; Duveau and Felix 2012; Andersen *et al.* 2014). While other QTL mapping studies of *C. elegans* behavior variation did not reach the QTN level, they sometimes revealed epistatic and interacting QTL in behavioral phenotypes (Gaertner *et al.* 2012; Bahrami and Zhang 2013; Glater *et al.* 2014) and, more generally, complex traits may often be influenced by multiple antagonistic-effect loci in tight linkage (Shao *et al.* 2008; Gaertner *et al.* 2012; Bernstein *et al.* 2018). Plasticity QTL for gene expression in *C. elegans* animals that had been reared at cold and hot temperatures reported a prominent role for trans-acting regulators, including a ‘master regulator’ with widespread effects (Li *et al.* 2006). Chronic exposure to distinct thermal regimes during development of distinct genotypes of *C. briggsae* reveals extensive modularity of plastic and genotype-dependent transcriptome responses (Mark *et al.* 2019), as do evolutionary responses to acute temperature stress in experimental populations of *C. remanei* (Sikkink *et al.* 2018). Should the temperature dependent behavioral QTL we uncovered provide a general source of temperature dependence of phenotypes and gene expression differences among genotypes, then it would point to genetically simple but highly pleiotropic underpinnings to adaptive differentiation between Temperate and Tropical populations of *C. briggsae*. The challenge remains to functionally characterize specific genetic variants controlling *C. briggsae*’s plastic thermal responses in behavior to test these additional aspects of its genetic architecture.

## Acknowledgements

We thank Joseph Ross for sharing genotype information for the RIL strains, Karl Broman for R/qtl troubleshooting and suggestions, Lanna Jin and Adrian Verster for help visualizing data in R. ADC was supported by funds from the Natural Sciences and Engineering Council of Canada and a Canada Research Chair. WSR was supported by funds from the Natural Sciences and Engineering Council of Canada.

